# FC2 stabilizes POR and suppresses ALA formation in the tetrapyrrole biosynthesis pathway

**DOI:** 10.1101/2022.10.12.511875

**Authors:** Tingting Fan, Lena Roling, Boris Hedtke, Bernhard Grimm

**Author notes:** both authors contributed equally to the work (Tingting Fan, Lena Roling). Corresponding author: Bernhard Grimm. Authors contribution: T.F. and B.G. designed the experiments. T.F., L.R. and B.H. performed experiments. T.F., L.R., B.H. and B.G. analyzed the data. B.G. wrote the manuscript. T.F., L.R. and B.H. prepared figures and contributed to the revision of the manuscript.

## Abstract

- During photoperiodic growth, the light-dependent nature of chlorophyll synthesis in angiosperms necessitates robust control of the production of 5-aminolevulinic acid (ALA), the rate-limiting step in the initial stage of tetrapyrrole biosynthesis (TBS). We are interested in dissecting the post-translational control of this process, which suppresses ALA synthesis for chlorophyll synthesis in dark-grown plants.
- Using biochemical approaches for analysis of wild-type and mutant lines as well as complementation lines, we show that the heme-synthesizing ferrochelatase 2 (FC2) interacts with protochlorophyllide oxidoreductase and the regulator FLU which both promote the feedback-controlled suppression of ALA synthesis by inactivation of glutamyl-tRNA reductase, thus preventing excessive accumulation of potentially deleterious tetrapyrrole intermediates.
- Thereby FC2 stabilizes POR by physical interaction. When the interaction between FC2 and POR is perturbed, suppression of ALA synthesis is attenuated and photoreactive protochlorophyllide accumulates. FC2 is anchored in the thylakoid membrane via its membrane-spanning CAB (chlorophyll-a-binding) domain.
- FC2 is one of the two isoforms of ferrochelatase catalyzing the last step of heme synthesis. Although FC2 belongs to the heme-synthesizing branch of TBS, its interaction with POR potentiates the effects of the GluTR-inactivation complex on the chlorophyll-synthesizing branch, and ensures reciprocal control of chlorophyll and heme synthesis.

## INTRODUCTION

Plants synthesize a broad range of tetrapyrroles, including chlorophyll (Chl), protoheme, siroheme and phytochromobiline, required for many essential cellular metabolic and signaling functions (Tanaka and Tanaka, 2007; Grimm, 2019). The photoreactive properties of Chl and tetrapyrrole intermediates necessitate tight control of tetrapyrrole biosynthesis (TBS) to ensure adequate flux of metabolites through the pathway and formation of the appropriate combination of end products, as absorption of light by free tetrapyrroles readily generates singlet oxygen. The rate-limiting step in TBS is catalyzed by glutamyl-tRNA reductase (GluTR), which performs the first step in the synthesis of 5-aminolevulinic acid (ALA). The activity, stability and subcellular localization of GluTR are controlled by multiple factors that serve to modulate the supply and allocation of ALA, which is the metabolic precursor of all tetrapyrroles (Czarnecki et al., 2011a; Wang et al, 2017; Meskauskiene et al., 2004; Richter et al., 2019). Plant tissues require varying quantities of heme and Chl, depending on the developmental stage and the environmental conditions considered. Therefore, the branch point in TBS, at which the appropriate amounts of protoporphyrin (Proto) are directed into either the Chl- or the heme-synthesizing pathways, must also be rigorously controlled. Mg chelatase (MgCh) and ferrochelatase (FC) insert Mg^2+^ and Fe^2+^ into Proto to produce Mg protoporphyrin (MgProto) and protoheme, respectively.

Finally, Chl biosynthesis in angiosperms necessitates rigorous control of light-dependent protochlorophyllide oxidoreductase (POR). Strict suppression of ALA synthesis is needed to avoid excessive accumulation of POR’s photoreactive substrate protochlorophyllide (PChlide) in the dark. This task is performed by the negative regulator FLU (FLUORESCENT IN BLUE LIGHT; Meskauskiene et al., 2004). FLU inactivates GluTR by physical interaction and forms a membrane-associated regulatory complex with POR, Mg protoporphyrin monomethylester (MgPME) cyclase (CHL27) and most likely other proteins (Kauss et al., 2012; Kong et al., 2016). It is assumed that the dark suppression of ALA synthesis is triggered by POR-bound Pchlide, which accumulates at night as a result of the blockade of POR activity (Gosling et al., 2004; Schmied et al., 2018).

Like all higher plants, Arabidopsis has two FC isoforms, FC1 and FC2, which share 83% similar amino acid residues. Low levels of FC1 mRNA continuously accumulate in all plant tissues, while FC2 is predominantly expressed in photosynthetically active leaf cells (Chow et al., 1998; Singh et al., 2002; Suzuki et al., 2002; Masuda et al., 2003). Relative to FC1, FC2 possesses an additional CAB (Chl-a/b-binding) domain, which facilitates binding of Chl, as has been shown for the cyanobacterial FC2-like homolog (Sobotka et al., 2011). The FC1 gene is strongly induced by various abiotic stress stimuli (Smith et al., 1994; Chow et al.; 1998; Singh et al., 2002; Nagai et al., 2007) and is, accordingly, responsible for induced tolerance to stress (Singh et al., 2002; Mohanty et al., 2006; Nagai et al., 2007; Scharfenberg et al., 2015; Zhao et al., 2017; Fan et al., 2019). FC2 was found to be exclusively located in chloroplasts, while FC1 was targeted to both plastids and mitochondria in organellar uptake experiments (Chow et al., 1997; Chow et al., 1998). The latter finding was subsequently questioned, when import of FC1 into Arabidopsis mitochondria failed (Lister et al., 2001), although later experiments detected FC activity in tobacco mitochondria, thus supporting the dual-targeting hypothesis (Hey et al., 2016). Only one chloroplast-localized FC isoform exists in Chlamydomonas reinhardtii, which displays greater similarity to the plant FC2 than to the FC1 isoform (van Lis et al., 2005). In contrast, the red alga Cyanidioschyzon merolae contains a mitochondrion-localized FC (Watanabe et al., 2013). The unresolved issue of the subcellular destination of the FC isoforms is reminiscent of the disputed organellar assignment of protoporphyrinogen oxidase (PPO), the TBS enzyme that acts upstream of FC. Like FC, PPO is encoded by two genes in all land plants. PPO1 is the dominant enzyme and is exclusively localized in plastids, while PPO2 has been reported to be targeted to mitochondria and/or chloroplasts in different angiosperm species (Watanabe et al, 2001; Lermontova et al., 1997; Chee et al., 2000).

Our central objective in this study was to assign specific functions in TBS to each FC isoform. FC1 has been shown to be indispensable for plant development and stress responses. Thus, FC1 knockout mutants of Arabidopsis are embryo-lethal, indicating that FC2 expression cannot compensate for the loss of FC1 (Fan et al., 2019). When FC2 was expressed under the control of the FC1 promoter, it successfully complemented the fc1 phenotype during embryogenesis, but failed to fully substitute for FC1 under stress conditions (Fan et al., 2019). Seedlings of FC2 knockout mutants show a necrotic phenotype and growth retardation owing to the accumulation of photoreactive Proto (Woodson et al., 2011). These results indicate that, in addition to their differential expression, the FC-encoding genes serve specific purposes, and it has been proposed that the two isoforms provide heme for varying sets of heme-dependent proteins (Singh et al., 2002; Zhao et al., 2017; Fan et al., 2019).

Heme is an indispensable cofactor for multiple fundamental biological processes. It is required for redox reactions in electron transport chains and other biochemical reactions, binds oxygen and other gases, acts as a cofactor for enzymes involved in responses to oxidative stress and for detoxification, such as peroxidases, catalases and cytochrome P450 proteins, and is involved in signaling processes and transcriptional control (Brzezowski et al., 2015; Balk and Schaedler, 2014; Larkin, 2016). FC1-overexpressing lines were first designated as genomes uncoupled 6 (gun6) mutants, because they partially restored expression of photosynthesis-associated nuclear genes (PhANGs) when chloroplast development was impaired, e.g., by Norflurazon treatment (Woodson et al., 2011). These findings support a regulatory role of FC1-synthesized heme in retrograde signaling during early chloroplast development (Woodson et al., 2011; Larkin, 2016; Chan et al., 2016). In contrast, overexpression of FC2 fails to evoke a gun-like phenotype (Woodson et al., 2011). In addition to intracellular signaling-dependent transcriptional control, heme also functions in post-translational control. Binding of heme to the GluTR-binding protein (GBP) attenuates its association with GluTR. As a result, GluTR becomes accessible to degradation by the Clp protease system, and ALA synthesis is diminished (Richter et al., 2019).

This experimental evidence for heme-mediated regulation in plants has prompted further studies on the impact of the two FC isoforms and heme in the control of TBS. In continuation of our previous work on the specific role of FC1 (Fan et al., 2019), we set out to examine the chloroplast-specific functions of FC2 that are not compensated for by FC1 in Arabidopsis FC2 knockout lines. As a result of the studies presented here, we identify FC2 as an additional factor that contributes to the feedback control of ALA synthesis at the level of the POR- and FLU-containing GluTR-inactivation complex. The association of FC2 with this complex stabilizes POR and ensures the attenuation of metabolic flow into Chl synthesis. It is proposed that the interaction of FC2 with the GluTR-inactivation complex not only reduces the supply of ALA for Chl synthesis, but also controls the production of heme for plastid-localized heme-dependent proteins.

## MATERIAL AND METHODS

### Plant materials and growth conditions

*Arabidopsis thaliana* seedlings were grown on soil under SD (8h light/16h dark cycles) or CL with 100 µmol photons m^−2^s^−1^ light intensity. Besides Columbia-0 (Col-0) as wild type, the T-DNA insertion mutants (*fc2-1* (GK_766_H08), *fc2-2* (SAIL_20_C06), *porb* (SALK_06019) and *flu*) have been described previously (Scharfenberg et al., 2015; Hey et al., 2016; Hou et al., 2019). For the generation of pFC2FC2(*fc2/fc2)* lines, the amplified genomic *FC2* sequence was inserted into pCAMBIA3301. For pFC2FC1(fc2/fc2) lines, the amplified *FC2* promoter sequence and the promoter-free genomic *FC1* sequence were ligated and inserted into pCAMBIA3301. The recombinant vectors were transformed into *fc2-2* heterozygous recipient plants. For 35SFC2 overexpression lines, the promoter-free genomic FC2 sequence was inserted into the linearized pGL1 prior to Col-0 transformation. PCR primers used are listed in Supplemental Table S1.

### Nucleic acid analysis

Genomic DNA extraction from Arabidopsis leaves was conducted as described previously (Fan et al., 2019). RNA was isolated with TriSure reagent (Bioline) according to the manufacturer’s instructions. The integrity of RNA was examined by gel electrophoresis, and the concentration determined with the NanoDrop 2000 Spectrophotometer (Thermo Scientific). One µg aliquots of RNA were digested with DNaseI (Thermo Scientific), and reverse transcribed by using RevertAid reverse transcriptase (Thermo Scientific). The qRT-PCR was performed with SYBR Green PCR master mix (Biotool) using the CFX96 real-time system (Bio-Rad). Gene expression was calculated by the 2^−ΔΔCt^ method and normalized to the reference gene *ACTIN*. qRT-PCR primers used are listed in Supplemental Table 1.

### Protein extraction and immunoblot analysis

Three-week-old Arabidopsis leaves were weighted, homogenized in liquid nitrogen and dissolved in 2x SDS sample buffer (100mM TRIS/HCl pH 6.8, 4% SDS, 20% glycerol, and 2mM dithiothreitol) at 95°C for 10min. For immunoblot analysis, equal amounts of proteins were separated on a 10% or 12% SDS-polyacrylamide (PA) gel and transferred to nitrocellulose membranes (GE Healthcare). The signal was probed with specific antibodies using Clarity Western ECL Blotting Substrate (Bio-Rad).

### Determination of tetrapyrroles and ALA synthesis rates

Tetrapyrroles were analyzed as previously described (Czarnecki et al., 2011a; 2011b). Leaf tissues were homogenized with liquid nitrogen and precursors and Chl were extracted in alkaline acetone (acetone:0.2N NH_4_OH, 9:1) at 4°C. After centrifugation, the pellet was resuspended with AHD buffer (acetone: dimethyl sulfoxide: 37% HCl, 100:20:5) to isolate noncovalently bound heme. The extracts were eventually separated and determined by high-performance liquid chromatography (HPLC) as described (Czarnecki et al., 2011b). ALA synthesis rates were measured according to (Mauzerall and Granick, 1956), with modifications (Richter et al., 2019).

### Analysis of thylakoid complexes

Thylakoid complexes were extracted and separated on a BN-PAGE gel according to (Jarvi et al., 2011), with modifications. For immunoblot analysis, the 1D-lane of separated proteins was sliced, denatured in SDS sample buffer for 30min at room temperature and loaded on a 10% or 12% SDS-PA gel containing 6M urea. After electrophoresis, the proteins were transferred to nitrocellulose membranes (GE Healthcare) for immune analysis with specific antibodies.

### Analysis of subplastidal localization

Four-week-old Arabidopsis plants were homogenized in buffer 1 (450mM Sorbitol, 20mM TRICINE, 10mM EDTA, 10mM NaHCO3, 0.1% BSA, pH 8.4) using a Waring blender. The suspension was filtered through Miracloth (Merck) and centrifuged for 8 min at 500xg. The pellet was resuspended in buffer 2 (300mM Sorbitol, 20mM TRICINE, 2.5mM EDTA, 5mM MgCl2, pH 8.4) using a soft brush and loaded on a step gradient of 40% and 80% Percoll (GE Healthcare) in buffer 2. Following centrifugation at 6500xg for 30 min, intact chloroplasts were washed using buffer 2 and pelleted for 6 min at 3800xg. Chloroplast were lysed osmotically in buffer 3 (25mM HEPES-KOH, pH 8.0) containing cOmplete protease inhibitor (Roche) and either separated into soluble and pellet fraction by centrifugation for 5 min. at 20,000xg or subjected to a separation of plastidal envelopes and thylakoids according to (Flores-Pérez and Jarvis, 2017).

### Pull-down assay

Ni-NTA agarose beads (Qiagen) were equilibrated in PBS buffer at 4°C. More than 40μg purified PORB-6xHis protein was ligated to Ni-NTA agarose by incubating at 4°C for 1h. Crude chloroplast extracts were solubilized in 1% (w/v) DDM on ice for 10min. The supernatant obtained after the solubilization was incubated with Ni-NTA agarose immobilized with the His-tag protein overnight at 4°C. As a negative control, the empty Ni-NTA agarose beads reacted with solubilized plastid extracts under the same condition. Subsequently, the agarose beads were washed at least three times with PBS buffer. The bound proteins were finally eluted and denatured in 2xSDS-PAGE sample buffer for immunoblot analysis.

### Bimolecular fluorescence complementation (BiFC) assay

This assay was performed according to a Venus YFP fluorescence system supplied with a GATEWAY vector construction system (Gehl et al., 2009). The final fluorescence was recorded with a Leica confocal laser-scanning microscope.

### Yeast two hybrid

The pDHB1(MCS2) and met25pXCgate (pNub) (46) vectors were applied as prey and bait constructs. The pDHB1-containing target gene was constructed by subcloning and restriction digestion of PCR-amplified sequences. The recombinant pNub vector was constructed by Gateway cloning via pDONR221. The transformation of yeast was conducted with modifications via a LiAc/SS carrier DNA/PEG method (Gietz and Schiestl, 2007).

## RESULTS

### FC2 Deficiency Impairs Chl Biosynthesis

The seedlings of the allelic mutants *fc2-1* (knock-down) and *fc2-2* (knock-out) show progressively slower growth rates, pale green pigmentation, and necrotic leaves in comparison to wild type (Fig. 1a-c). The similar phenotype has been described for these mutants in previous reports (Scharfenberg et al., 2015; Woodson et al., 2015; Espinas et al., 2016). The severity of the phenotype differs between the two mutants (Fig. 1d). While *fc2-2* is devoid of FC2 and grows only in continuous light (CL), the low FC2 level in fc2-1 permits growth of seedlings under both short-day (SD) and CL conditions, but the mutant leaves are pale green and develop necrotic spots. Besides confirming previous findings for the FC2-deficient mutants, we analyzed the steady-state amounts of their tetrapyrrole end products and intermediates as well as the levels of representative TBS proteins (Fig. 2). Chl and non-covalently bound (ncb) heme contents were decreased in both SD and CL-grown *fc2-1* and *fc2-2*. In FC2-deficient plants, reduced amounts of proteins involved in Chl synthesis were observed, while the content of GluTR and GSAAT involved in ALA synthesis remained stable. Compared to wild type, the level of the heme-binding cytochrome *f*, but also POR and the MgCh proteins CHLH and GUN4 of the Chl-synthesizing branch were remarkably lower in the *fc2* mutants. As result of lower Chl content, the FC2-deficient seedlings also displayed a concomitant loss of the antenna and core proteins of the photosystems (Figs. 2c and S1).

**Fig. 1.**
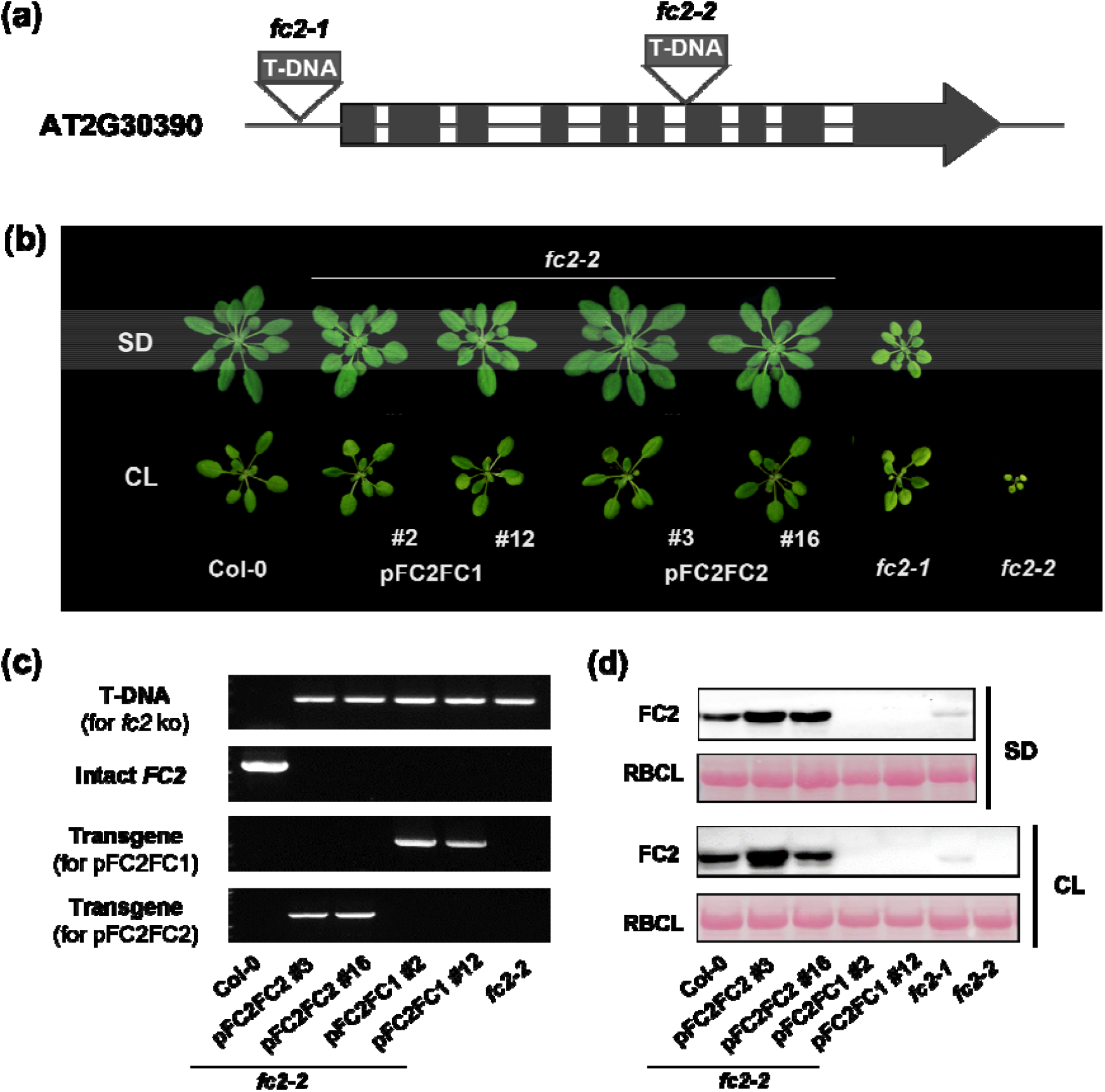
Phenotypic characterization of *fc2-2-*complemented plants expressing *FC1* or *FC2* under the control of the *FC2* promoter. (**a)** Schematic depictions of the *FC2* (AT2G30390) locus with two allelic T-DNA insertion mutants. Introns (white boxes) and exons (black boxes) are indicated. (**b)** Comparison of seedlings from pFC2FC1(*fc2/fc2)* and pFC2FC2(*fc2/fc2)* complementation lines with wild-type and *fc2* mutant plants. Plants were grown under short-day (SD) conditions for 5 weeks or in constant light (CL) for 2 weeks. (**c)** Genotyping analyses of pFC2FC1(*fc2/fc2*), pFC2FC2(*fc2/fc2*) and *fc2-2* mutants, as well as wild-type seedlings. **(d)** Immunoblot analysis of FC2 protein contents in leaves of 5-week-old seedlings of wild-type, pFC2FC1(*fc2/fc2*), pFC2FC2(*fc2/fc2*), *fc2-2* and *fc2-1* lines grown under SD or CL conditions

**Fig. 2.**
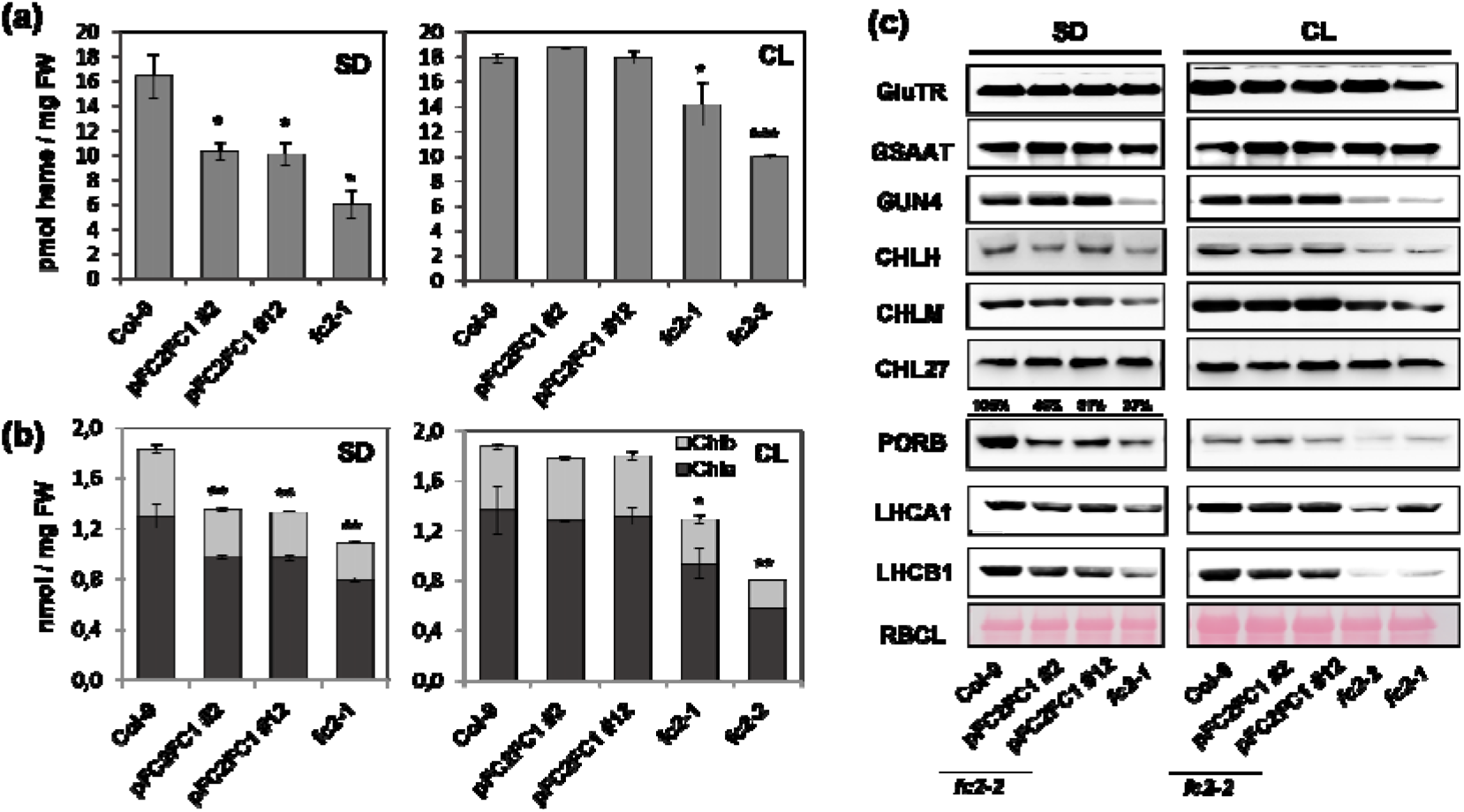
Biochemical and genetic analysis of complemented *fc2-2* lines (pFC2FC1(*fc2/fc2))* and wild-type controls. **(a**,**b)** Quantification of end products of TBS. Heme (A) and chlorophyll (B) were extracted from leaf samples harvested from 3-week-old SD-grown and 2-week-old CL-grown seedlings of pFC2FC1(*fc2/fc2*), *fc2* and wild-type plants. (c) Analyses of selected proteins in SD- and CL-grown plants. GluTR: Glutamyl-tRNA reductase; GSAAT: Glutamate-1-semialdehyde aminotransferase; FLU: Fluorescent; GBP: GluTR binding protein; CHLH: Mg chelatase subunit H; GUN4: genomes uncoupled 4; CHLM: Mg-protoporphyrin IX methyltransferase; CHL 27, Mg protoporphyrin monomethylester (MgPME) cyclase; POR B: protochlorophyllide oxidoreductases B; LHCA1/LHCB1: Light-harvesting chlorophyll-binding protein A1 of photosystem I and B1 of photosystem II; RBCL: ribulose 1,5-bisphosphate carboxylase oxygenase, large subunit.

Biochemical and genetic analysis of *fc2-2* seedlings transformed with *pFC2:FC2* revealed a complete phenotypic rescue (Fig. 1b) and wild-type contents of TBS proteins and end products were restored (Fig. S2). In contrast, expression of *FC1* under the control of the *FC2* promoter *(pFC2:FC1)* resulted in partially complemented *fc2-2* lines. SD-grown pFC2FC1(*fc2/fc2)* seedlings were slightly pale green and smaller than those of the control lines (Fig. 1b). The lack of full complementation was substantiated by the analysis of ncb heme and Chl contents in light-exposed leaves. SD-grown pFC2FC1(*fc2/fc2*) lines accumulated 27% and 38% less Chl and ncb heme, respectively, than wild-type control (Fig. 2a-b).

Interestingly, among the analyzed TBS proteins lower levels of PORB were not only detected in the *FC2-*deficient lines under CL and SD conditions, but also in pFC2FC1*(fc2/fc2) lines* under SD. The other representative proteins of the Chl synthesis branch accumulated to wild-type levels (Fig. 2c).

### FC2 is Important for the Stabilization of the PORB-FLU-GluTR Complex and Control of ALA synthesis

To further explore the diverse functions of the two FC isoforms and to detect reasons for the only partial complementation of *fc2* with *FC2*-promoter-driven *FC1* expression, we assayed protein extracts from two representative pFC2FC1 lines lacking FC2, *fc2-1* and two *FC2* overexpressor lines and compared these plants with wild type (Fig. 3a-b). We included to this analysis the *porb* knock-out mutant, as the PORB content was initially found to be strikingly diminished in FC2-deficient lines (Fig. 2c). This was confirmed and, interestingly, FC2-overexpressing lines contained elevated POR contents (Fig.3b). Other TBS proteins, particularly those involved in ALA synthesis, accumulated to similar amounts in all analyzed lines. These results are compatible with FC2-dependent stabilization of PORB. The observed interdependency of heme-synthesizing FC2 with PORB contents points to an association to the potential Chl accumulation.

**Fig. 3.**
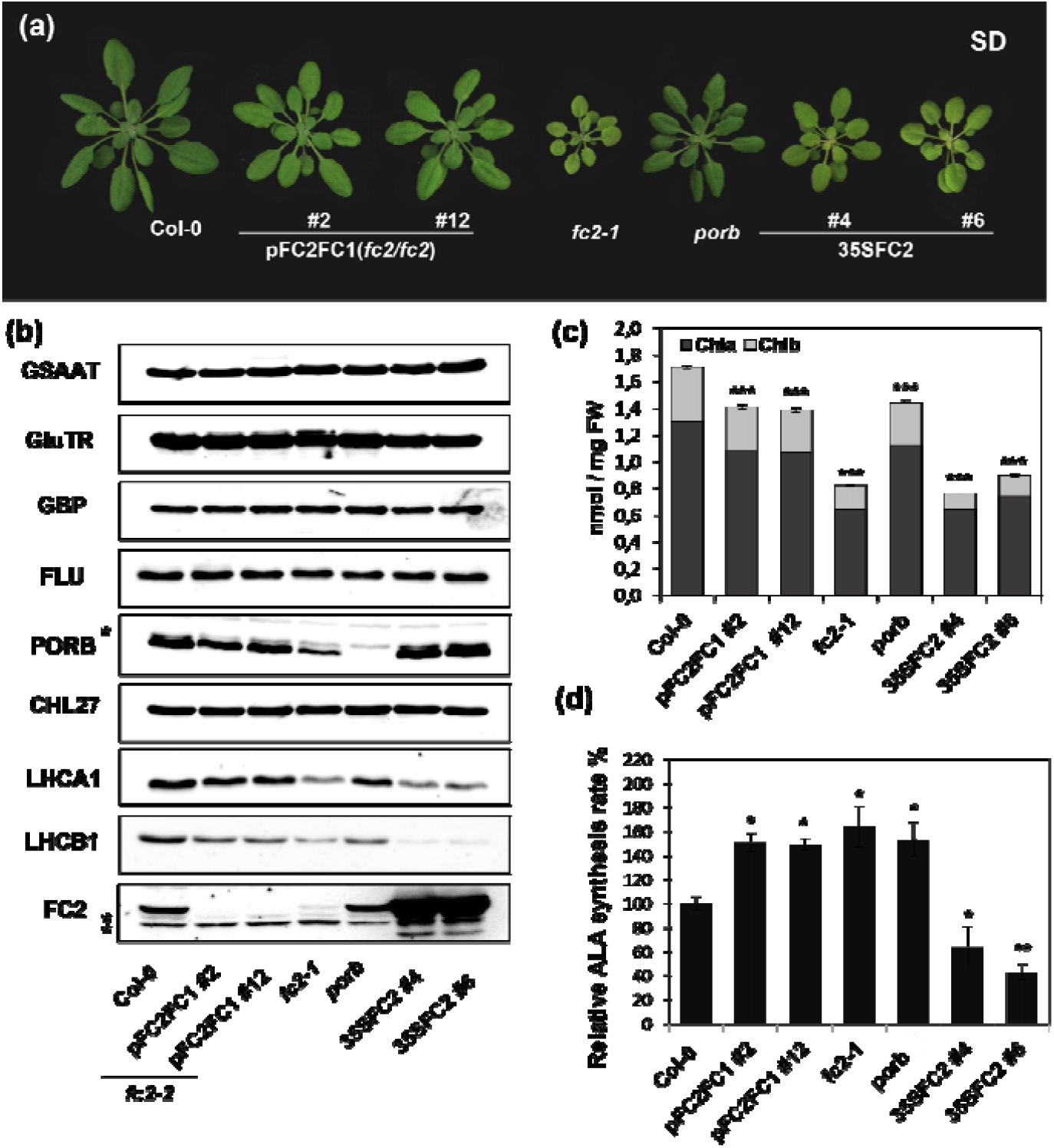
Characterization of *FC2* overexpression lines and porb in comparison to pFC2FC1(*fc2/fc2*), *fc2-1* and wild type under SD conditions. **(a)** Phenotypes of 5-week-old porb mutants and 35SFC2 overexpression plants grown under SD conditions. (b) Western blot analysis of TBS proteins and LHCPs in SD-grown mutants and wild-type seedlings. Asterisks indicate non-specific immunoreactive bands. **(c**,**d)** Determination of chlorophyll (c) and relative ALA synthesis rates (wild type corresponds to 100 pmol ALA h^-1^ g FW^-1^) (**d)** in 3-week-old mutants and comparable wild-type seedlings from panel (**a**). Error bars indicate standard deviations (n≥3). Asterisks represent significant differences, as determined by Student’s t-test; * *P* < 0.05, ** *P* < 0.01, *** *P* < 0.001.

The *FC2* overexpressor lines contained only half as much Chl as the wild type, equivalent to the amount detected in *fc2-1*. In marked contrast, Chl accumulation in pFC2FC1(*fc2/fc2)* lines was only slightly lower than in wild type and similar to that in *porb* (Fig. 3c). As expected, the reduced Chl contents in the *fc2-1* and FC2-overproducing lines were accompanied by decreased level of Chl-binding proteins, exemplified by LHCB1 and LHCA1 (Fig. 3b).

We further compared ALA synthesis rates and steady-state levels of PChlide in the mutants with different levels of FC2 and PORB relative to wild type. Interestingly, the ALA synthesis rate was higher in all FC2-deficient lines as well as in porb and lower in the FC2 overexpressing lines indicating that FC2 and PORB contents were inversely correlated with the ALA synthesis rate (Fig. 3d). As the total contents of GluTR, GSAAT, GBP as well as FLU were unchanged in all transgenic lines, the results suggest that FC2 posttranslationally modulates ALA synthesis rates.

### FC2-PORB Interaction and the Association of GluTR with the Thylakoid Membrane

ALA synthesis is controlled by several mechanisms. Two regulatory factors dominantly affect the stability and activity of GluTR. Firstly, GBP stabilizes the ALA-synthesizing complex (Sinha et al., 2022). Apart from that, binding of heme to GBP weakens its interaction to GluTR in the stroma and thereby makes GluTR accessible to proteolytic degradation by the Clp protease (Richter et al., 2018). Secondly, during darkness membrane-bound FLU inactivates the large part of GluTR in the inactivation complex in response to Pchlide-associated POR (Meskauskine et al., 2010). Both a deficit and an excess of membrane-localized FLU were previously reported to change the distribution of GluTR between the thylakoid membrane and the stroma of seedlings grown under light-dark cycles (Hou et al., 2019). In this way, the different subplastidal localizations of GluTR directly affect the ALA synthesis rate. An inverse correlation between the amount of membrane-bound GluTR and the rate of ALA synthesis has been reported previously (Schmied et al., 2018).

The impact of the FC2 level on the distribution of GluTR between the chloroplast stroma and membranes was therefore determined in seedlings of *fc2-2*, pFC1FC2*(fc2/fc2)*, an FC2-overexpressing line, *porb* and wild type (Fig. 4a). While GluTR amounts associated with the thylakoid membranes increased after the transition from light to dark in wild type, dark-grown FC2-deficient lines displayed reduced amounts of membrane-bound GluTR. Conversely, membrane-associated GluTR contents were elevated in the p35SFC2 line relative to those in light-exposed leaf samples of these lines.

**Fig. 4.**
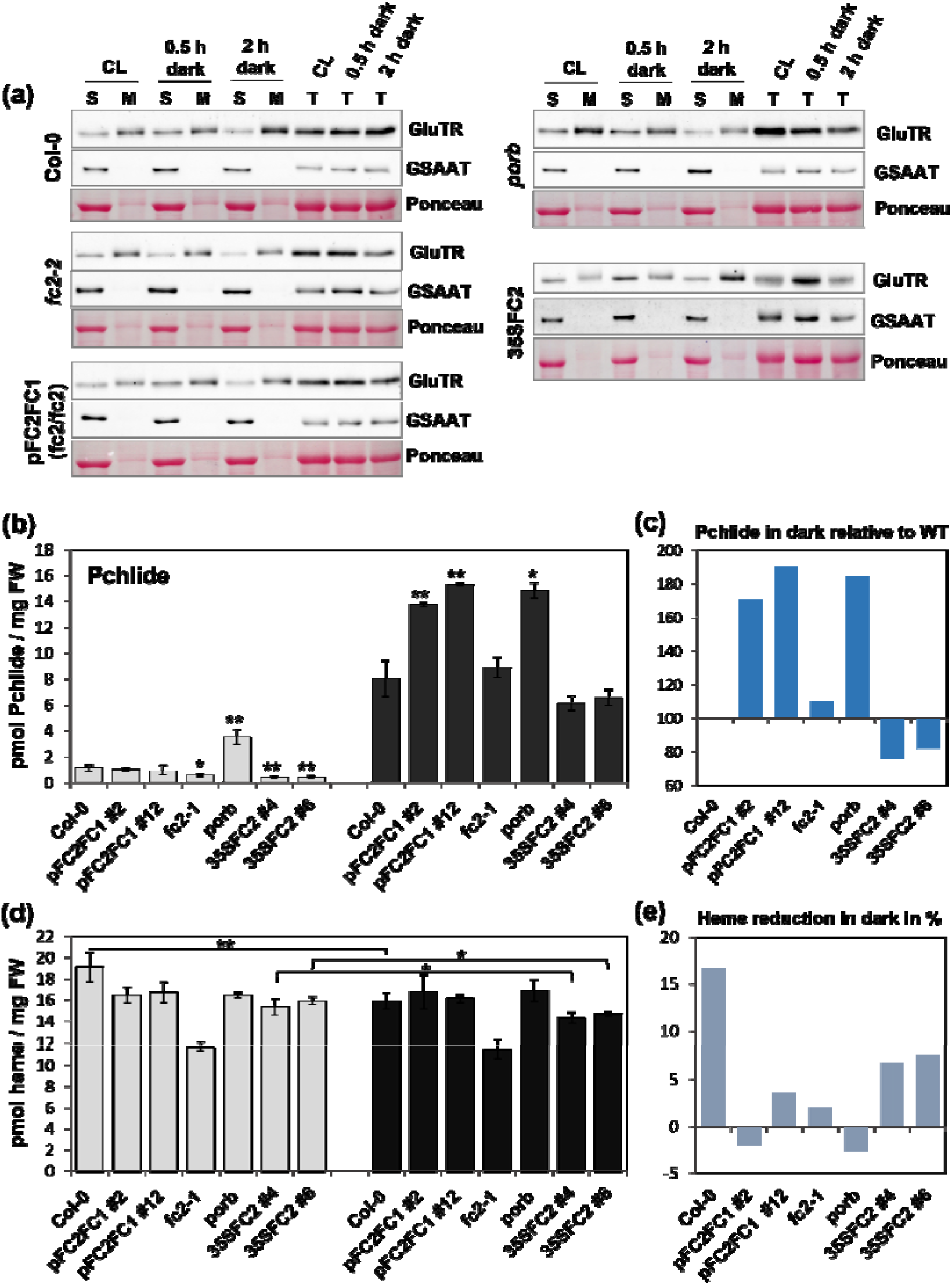
Deficiency of FC2 or PORB affects GluTR localization as well as heme and Pchlide accumulation in darkness. **(a)** GluTR distributions between membrane and stroma fractions. Leaf samples of wild type and mutants were separated into soluble and pellet fractions, which represent proteins of the plastid stroma (S) and thylakoid membranes (M), respectively. Leaves were harvested from 4-week-old, CL-grown seedlings prior to (CL), and after a period of 30 min (0.5 h) or 2 hours (2 h) of darkness. **(b**,**d**) Accumulation of Pchlide (**b**) and heme **(d)** in CL (white boxes) and dark-incubated (white boxes) plants. Dark samples were harvested after 12h in darkness. Error bars indicate standard deviations (n≥3). Asterisks represent significant differences, which were calculated with reference to light- and dark-grown Col-0, respectively; * *P* < 0.05, ** *P* < 0.01. *P* values of heme content were calculated by comparing light- and dark-grown samples. (**c)** Changes in Pchlide accumulation in darkness, relative to wild type. Increase in Pchlide content of dark-exposed Col-0 was set to 100%. **(e)** Heme reduction in darkness, relative to content in light (in %). Changes were calculated by comparing heme contents in light- and dark-exposed plants. Heme contents of light-grown samples were set to 100%

Our results indicated an inverse correlation between levels of FC2/PORB and ALA synthesis, which is affected by the distribution of GluTR between stroma and membrane (Figs. 3b,d and 4a). It is therefore proposed that a lower FC2 content leads to a decrease in the GluTR content on the membrane, which correlates with less effective suppression of ALA synthesis. Conversely, higher levels of FC2 promote membrane-binding of GluTR, thus enhancing dark-inactivation of ALA synthesis.

Further evidence for a FC2-dependent impact on the efficacy of the GluTR inactivation complex was obtained by comparing levels of PChlide and heme levels in mutants with modified FC2 contents in light and darkness with those in wild type (Figure 4b-e). In light steady-state levels of Pchlide remain very low, since Pchlide is instantaneously converted to Chlide by POR. Elevated Pchlide content in the *porb* mutant points to an insufficient compensatory activity of the other two POR isoforms. At night, PChlide accumulated to several-fold higher levels in wild-type seedlings than during light exposure. Interestingly, FC2 overexpression correlated with decreased Pchlide levels both in light and darkness, which could be explained by enhanced GluTR inactivation and lower ALA synthesis rates (Figure 4b). In contrast, the dark-grown pFC2FC1(*fc2/fc2*) and *porb* seedlings accumulated about 180% of wild-type Pchlide content. These modified Pchlide levels point out to an impaired dark-suppression of ALA synthesis. Under the same conditions, *fc2-1* displayed decreased Pchlide accumulation in daytime compared to the control line, but a wild-type level during the night. In conclusion, the elevated Pchlide levels during the dark period correlate with an insufficient repression of dark ALA synthesis and, in contrast, enhanced suppression of ALA synthesis in FC2-overproducing lines parallels a lower accumulation of Pchlide in darkness.

In addition to evaluating the impact of FC2 accumulation on the flow of tetrapyrrole metabolites into the Chl synthesis branch, we determined its effect on heme content during light and dark incubation. Previous reports point to scarce differences in content of measurable heme between day and nighttime, but to mutants with impaired TBS displaying always reduced heme levels (Czarnecki et al., 2011a; Alawady and Grimm, 2005; Lermontova and Grimm, 2006; Apitz et al., 2016). As a consequence of down-regulated ALA synthesis in darkness, a 15% drop in ncb heme content was observed in wild-type seedlings after a 12h dark period (Fig. 4d-e). The heme levels in the two *35SFC2* lines were always lower than in the wild-type control, which correlates with lower ALA synthesis rates. These lines accumulated 7% less ncb heme in the dark compared to the light-exposed plants. The *fc2-1* seedlings contained only 61% of wild-type heme content, which did not change overnight. The pFC2FC1(*fc2/fc2*) and *porb l*ines accumulated equal amounts of ncb heme during light and dark incubation and thus, showed no reduced dark heme content unlike wild type. In summary, heme levels in the FC2/PORB-deficient lines did not decrease during dark incubation relative to the values in light-exposed samples. This also argues for impaired dark suppression of ALA synthesis when either FC2 or PORB are missing (Fig. 3d).

### Localization of the FC Isoforms in Chloroplasts

Due to the only partial complementation of the *FC2* knockout mutant by *pFC2:FC1* expression in leaves and the impaired ALA synthesis rate and heme accumulation in light-as well as dark-grown FC2-deficient and -overproducing seedlings, we asked for the subplastidal localization of both FC isoforms. We assessed the subplastidal localization of the two FC isoforms in wild-type chloroplasts and in a representative FC1 overexpressor line. The latter transgenic line was chosen, because FC1 is hardly detectable in protein extracts of wild-type leaves. We assumed that overproduced FC1 can be assigned to the same plastid subcompartments as in wild type, although it cannot be entirely excluded that excess FC1 occupies FC2 binding sites, in particular in FC2-deficient seedlings. We divided total leaf protein extracts into soluble and membrane-associated fractions. The plastid membranes were then further fractionated using sucrose-gradient centrifugation to enrich for chloroplast envelope proteins. For comparison, we determined the localization of the two PPO isoforms in plastids.

FC2 was exclusively detected in the thylakoid membrane fraction, as was PPO1. A substantial portion of overproduced FC1 was detected in the stromal fraction. However, a significant amount of FC1 was also found in the envelope fraction, while a large portion of overexpressed FC1 accumulated in the thylakoid membrane. The presence of the envelope marker protein Tic110 in the thylakoid fraction suggests some contamination of the thylakoid fraction with envelope proteins (Fig. 5), which would result in an underestimation of envelope-bound FC1. The observed presence of FC1 in the stroma could be explained by its lack of a clearly hydrophobic transmembrane domain like the C-terminal CAB domain of FC2. The assignment of FC1 to both plastid membrane fractions hints at possible associations of FC1 with additional proteins bearing integral membrane domains. Interestingly, PPO2 was predominantly found in the envelope fraction. Similar to the different levels of the two FC isoforms in leaf cells, it is worth mentioning that PPO2 constitutes only a minor portion of the total PPO content.

**Fig. 5.**
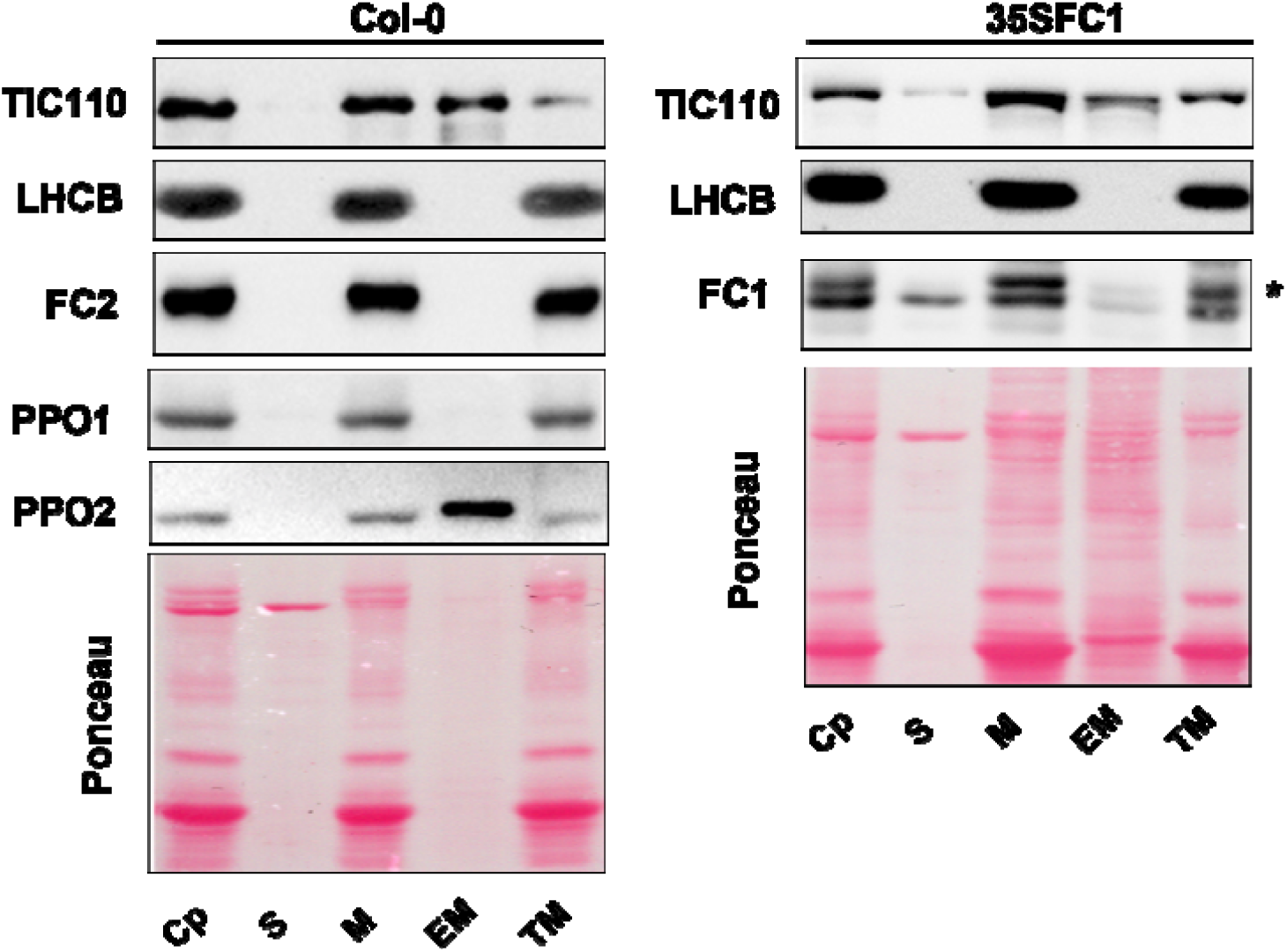
Suborganellar localization of FC and PPO proteins. Chloroplasts were isolated from Col-0 and 35SFC1 overexpressor seedlings. Lysed chloroplasts (Cp) were divided in soluble (S) and membrane (M) fractions. The latter were further fractionated by centrifugation on a three-step sucrose gradient to yield envelope membranes (EM) and thylakoid membrane (TM) fractions. Immunoblots were performed with antibodies specific for TIC110, LHCB1, FC1, FC2, PPO1 and PPO2. Ponceau-stained membranes are depicted; arrowheads highlight the large subunit of RuBisCO. The asterisk indicates a non-specific immuno-reactive band

### FC2 Interacts with All Three Isoforms of POR

The enhanced ALA synthesis rate and lower Chl content observed in *fc2-2* expressing pFC2:FC1 (Fig. 3d) and the correlation between accumulation of FC2 and the stability of PORB (Fig. 2c) prompted us to analyze a potential FC2–POR interaction. This interaction was verified by Bimolecular Fluorescence Complementation (BiFC) assays (Fig. 6a). Fig. S3 shows FC2 interaction with both PORA and PORC. FC2 interaction was also demonstrated with PPO1, the dominant plastidic isoform acting upstream of the Chl and heme-synthesizing branch (Fig. S4). The interactions of FC1 and FC2 with all three POR isoforms were analyzed using the yeast-two-hybrid (Y2H) approach (Fig. 6b). FC2 interacts with each of the three POR isoforms, while analysis of FC1 revealed no association with POR proteins. The Y2H experiment also confirmed the formation of FC2 homodimers, as well as heterodimers of FC2 and FC1. Expression of FC2ΔCAB, which lacks the coding sequence for the C-terminal domain, did not result in detectable interactions with POR (Fig. 6b). This CAB domain of FC2 constitutes the main structural difference between FC2 and FC1, functions as an anchor in the thylakoid membrane (Sobotka et al., 2005), and is proposed to represent a potential POR-binding site of FC2.

**Fig. 6.**
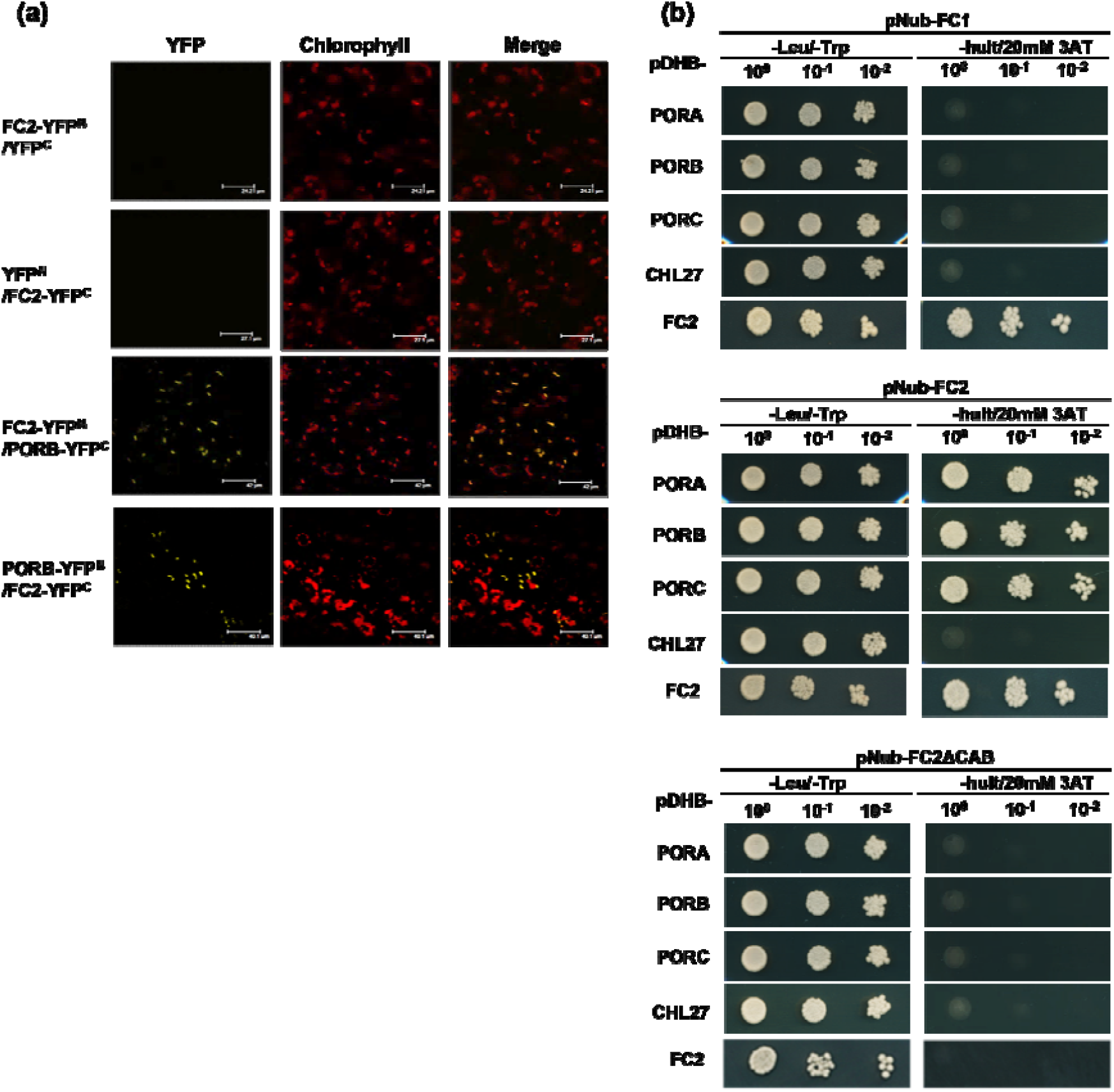
Analysis of FC2-PORB interaction. **(a)** BiFC analysis of interaction was performed by transiently expressing FC2 and PORB in leaf epidermal cells of *Nicotiana benthamian*a plants. Yellow fluorescence derives from the YFP tag; red fluorescence represents chloroplast autofluorescence; overlay images show merged signals. Both the combination of N-terminally (YFP^N^)-tagged FC2 and C-terminally (YFP^C^)-tagged PORB or the reverse pairing showed interaction in chloroplasts. Two negative controls expressing YFP^N^-tagged FC2 with YFP^C^ protein and YFP^C^-tagged FC2 with YFP^N^ protein generated no fluorescent YFP signals. **(b)** Interactions of FC2 and POR proteins in yeast cells. The yeast strain LC40 ccuα expressing pNub-FC1, pNub-FC2 and pNUB-FC2ΔCAB, respectively, was crossed with LC40 ccuA cells carrying pCub-PORa, pCub-PORB, pCub-PORC, pCub-CHL27 and pCub-FC2. Specific interactions were identified by the growth of transformants on SD/-Leu/-Trp/-His/-Urea (-hult) medium containing 20 mM 3-amino-1,2,4-triazole (3AT), a competitive inhibitor of the HIS3 enzyme (imidazole glycerol-phosphate dehydratase).

### FC2 Associates with the PORB-FLU-GluTR Complex

As FC2 and PORB act in different branches of TBS, their interaction suggests the existence of a common regulatory mechanism for heme and Chl synthesis. Because of the modified ALA synthesis and heme content of FC2-overexpressor lines as well as pFC1:FC2(*fc2/fc2*) in comparison to wild type, we searched for a common and specific role of FC2 and POR. We considered the GluTR-inactivation complex as a putative site of the FC2-POR interaction and assayed for additional FC2 connection with other proteins of TBS. Kauss et al. (2012) defined this membrane-localized protein complex in darkness as to consist at least of FLU, GluTR, POR and CHL27. FC2 showed no interaction with the MgPME cyclase subunit CHL27 (Fig. 6b, S3). However, interaction of FC2 with GluTR and FLU was detected in BiFC assays (Fig. 7a). As an additional positive control, BiFC assays confirmed PORB interaction with GluTR and FLU as well as the GluTR-FLU interaction.

**Fig. 7.**
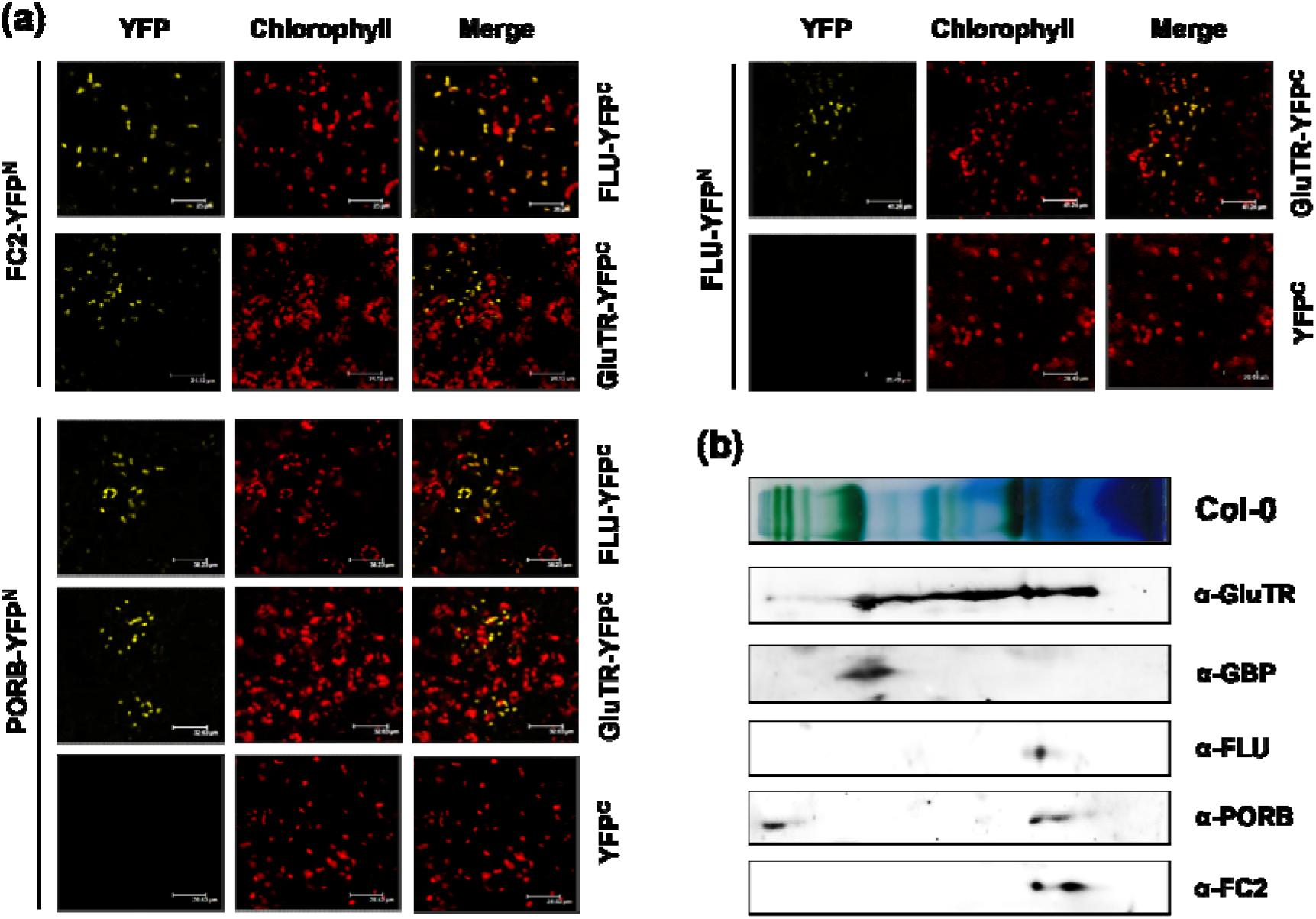
FC2 is associated with the POR-FLU-GluTR complex. **(a)** Visualization of protein interactions in chloroplasts using BiFC assays. YFP images depict reconstituted YFP fluorescence in tobacco cells transiently expressing constructs encoding the fusion proteins indicated at the left and right sides. Red fluorescence (Chlorophyll) represents chloroplast autofluorescence, and overlay images (Merge) show merged signals. Each image is representative of more than three replicate experiments. **(b)** Allocation of FC2, POR, GluTR and its regulatory proteins FLU and GBP to different protein complexes associated with thylakoid membranes. Solubilized wild-type thylakoids were separated on a BN-PAGE gel in the first dimension, followed by SDS-PAGE analysis in the second dimension. The proteins were immune-detected using the specific antibodies indicated.

Further experimental support for the interaction of FC2 with the PORB-FLU-GluTR complex was obtained by two-dimensional Blue Native (2D-BN) polyacrylamide gel electrophoresis (PAGE) of wild-type plastidal membrane extracts. Size fractionation in the second dimension of the BN PAGE revealed a comigration of FC2, PORB, FLU and GluTR (Fig. 7b). A second slightly smaller immune signal was also obtained for PORB and FC2. The immune signals for the four TBS enzymes correspond to protein complexes with an approximate molecular mass of less than 120.000 Dalton (Figure 7b). Comigrating immune-reacting spots of different proteins could point to a common protein complex. But due to the molecular mass of migrating protein complexes in the BN-gel, the corresponding complex cannot comprise all four proteins. It is not excluded that the leaf extraction protocol used does not allow maintaining a stable inactivation complex with these proteins.

Alternatively, it is suggested that the protein complexes of comparable size are dynamically assembled.

## DISCUSSION

The differential expression of *FC1* and *FC2* in Arabidopsis suggests that they have distinct functions, and, more specifically, that the two isoforms feed different heme pools and contribute to different regulatory mechanisms (Chow et al., 1998; Suzuki et al., 2002, Nagai et al., Woodson et al., 2015, Fan et al., 2019). *FC2* is the dominant isoform in photosynthetic leaf tissue, and its loss results in necrotic leaf lesions and seedling lethality during photoperiodic growth (Fig. 1, Scharfenberg et al., 2015). *FC2* antisense tobacco lines also display leaf necroses caused by Proto accumulation (Papenbrock et al., 2016). These observations support a predominant contribution of FC2 to heme synthesis in photosynthetic tissues. Conversely, *FC1* expression is essential for embryonic development and stronger in sink tissue, such as roots and flower organs, compared to leaves (Fan et al., 2019, Hey et al., 2016, Singh et al., 2002, Nagai et al., 2007). This differential expression was suggested to explain the embryo-lethality of a *FC1* knock-out mutant, as well as the wild-type-like appearance of *FC1* knock-down mutants under standard growth conditions (Espinas et al., 2016; Fan et al., 2019).

In light of the proposed divergent functions of the two isoforms of FC, we explored the capacity of FC1 to complement *fc2-2* by expressing FC1 under the control of the *FC2* promoter. While endogenous FC1 cannot substitute for FC2, as shown in *fc2-2* during photoperiodic growth (Scharfenberg et al., 2015; Woodson et al., 2015; Fig. 1), *FC2* promoter-driven expression of FC1 partially compensates for the lack of FC2 function (Fig. 1). Even overexpression of *FC1*, driven by the 35S promoter, failed to complement the loss of FC2 entirely (Woodson et al., 2015). Expression of *pFC2:FC1* in *fc2-2* mitigated the necrotic phenotype in continuous light, but not under light-dark conditions, confirming the importance of FC2 for photoperiodic growth. The incomplete complementation of *fc2-2* by *FC1* expression is reminiscent of the partial complementation of *fc1* achieved by *pFC1:FC2* expression, which was evidenced by an inadequate stress response (Fan et al., 2019). Similarly, expression of the transgene pFC2:FC2 did not rescue the FC1 deficiency during embryogenesis (Fan et al., 2019). The incomplete complementation of *fc1* and *fc2* by the alternative isoforms expressed under control of the specific promoters points to specific roles for each in the maintenance of plant heme synthesis.

### The distinct subcellular localization of the two FC isoforms precludes complete complementation of FC knock-out mutants

Our studies on the plastid localization of the two FC isoforms confirm that FC2 is found exclusively in thylakoid membranes, while FC1 was detected in the stromal fraction, bound to the envelope as well as attached to thylakoid membranes (Fig. 5). Because of low abundancy of FC1, the protein was not immunologically detectable in the subplastidal compartments of wild-type Arabidopsis, but only in *FC1-*overproducing lines. Based on this immune analysis, we assume that FC1 allocation covers several plastidal compartments (most likely in dependency to interacting proteins). Thus, we tentatively hypothesize that FC1 can be anchored by PPO2 to the chloroplast envelope, resulting in an association of FC1 with this membrane. The dominant isoforms PPO1 and FC2 were demonstrated to interact (Fig. S4) and are both exclusively located in the thylakoid membrane (Fig. 5), where Proto supplied by PPO1 is proportionally distributed into the heme and Chl-synthesizing branches (Fig. 8). While physical interaction of PPO1 with FC2 in the thylakoid membranes facilitates substrate channeling, the molecular interactions between PPO and MgCh and the delivery of Proto for MgCh await further studies. The MgCh complex was found to be attached to the thylakoid membranes (Gibson et al., 1996; Adhikari et al., 2009).

**Fig. 8.**
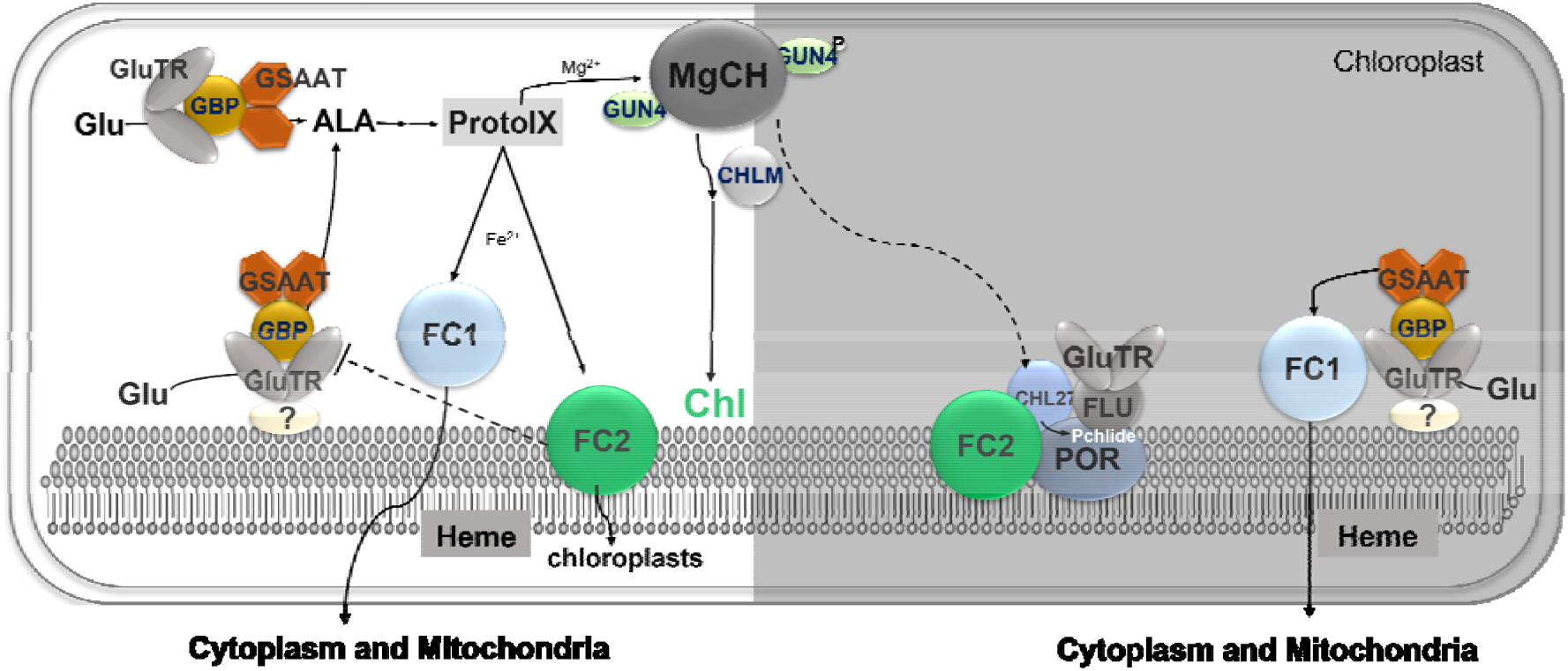
Working model. During the day (left side of scheme), chloroplasts efficiently synthesize ALA, thanks to an interaction between GSAAT and GluTR. The dimeric GSAAT binds to the V-shaped GluTR dimers, mainly in stroma, while a minor portion of GluTR is attached to the thylakoid membranes. During light exposure, GUN4 stimulates MgCh activity in the chlorophyll-synthesizing branch of TBS. Meanwhile, both FC isoforms catalyze the insertion of ferrous iron into Proto to yield heme. In the dark, FC2 interacts with and stabilizes the enzyme PORB (right side). As the activity of light-dependent POR is inhibited in the dark, Pchlide can accumulate and triggers the formation of a complex consisting of (at least) FLU, POR and CHL27. Formation of this regulatory complex enables inhibition of ALA synthesis by interaction of FLU with GluTR (GluTR-inactivation complex). Repression of ALA formation reduces the synthesis of both heme and chlorophyll in the dark, in order to coordinate the supply of both tetrapyrroles for plastid-localized components of the photosynthetic protein complexes, and provide for photoprotection upon illumination.

The robust integration of FC2 into the membrane can be readily explained by its membrane-spanning CAB domain, which facilitates specific anchoring of FC2 to thylakoids. Indeed, lack of the CAB domain also prevents the interaction of FC2 with POR (Fig. 6b). Previous studies had revealed that the CAB domain of cyanobacterial FC is essential for dimer formation, and plays a regulatory role in Chl synthesis and accumulation of Chl-binding proteins (Sobotka et al., 2011). Dimeric FC was shown to bind Chl between the CAB domains, while monomeric FC remains unpigmented (Pazdernik et al., 2019). It remains a challenging task to explore how the CAB domain of Arabidopsis FC2 determines the specific localization to the thylakoid membrane and interactions with other TBS proteins. Moreover, it is tempting to examine in the future whether the addition of FC2’s CAB domain to FC1 would enable the latter to fully substitute for FC2 itself.

Due to its high expression level in leaf cells and its localization to thylakoid membranes (Fig. 5) we can confirm that FC2 has a predominant role in the supply of heme for plastid-localized hemoproteins (Espinas et al., 2016). The different localizations of the two FC isoforms within the chloroplast imply that they perform different tasks and likely explain why light-exposed seedlings of FC2-deficient *fc2-1* and pFC2FC1(*fc2/fc2*) did not accumulate more Chl or heme, despite their elevated ALA synthesis capacity. It is assumed that the specific location of FC1 within the chloroplasts explains why it cannot completely compensate for loss of FC2.

### FC2 associates with the POR-FLU-GluTR inactivation complex

The physical interaction of FC2 with POR was demonstrated by BiFC and Y2H assays (Figs. 6 and S3). In contrast, FC1 did not interact with any of the three POR isoforms in Arabidopsis. These studies excluded a direct interaction of FC2 with CHL27. Furthermore, BiFC assays detected associations of GluTR and FLU with FC2, although an indirect interaction mediated by endogenous POR cannot be excluded (Fig. 7a). Fractionation of membrane-associated protein complexes by 2D BN-PAGE revealed consistent co-migration of FC2 with a subpopulation of POR, FLU and GluTR, all of which are known to form the GluTR-inactivation complex (Kauss et al., 2012; Fig. 7b). The two immunoreactive FC2 spots identified in the 2D-BN-PAGE experiment suggest that some FC2 is associated with POR, while the rest presumably represents the catalytically active FC2, which could interact with other proteins. Note that co-migration of these proteins was observed in leaf samples harvested after 30 min of dark incubation, which is sufficient to trigger dark-suppression of ALA synthesis (Richter et al., 2010).

The GluTR-inactivation complex serves to control ALA synthesis in the dark as previously demonstrated (Meskauskiene et al., 2001), but is also required during the day under fluctuating light levels and presumably other changing environmental conditions (Hou et al., 2019). Thus, it is assumed that the assembly of proteins into the inactivation complex is observable not only during darkness, but also as an additional post-translational control mechanism in the light.

### FC2 deficiency correlates with elevated ALA synthesis

The interaction of FC2 with the GluTR inactivation complex is compatible with known aspects of the rather complex control of ALA synthesis. Here, the regulatory influence of different forms of the inactivation complex was demonstrated in FC2-deficient and overproducing lines. Enhanced ALA synthesis rates were observed in FC2-deficient seedlings (*fc2-1* or pFC2FC1(*fc2/fc2*) lines), although GluTR contents remained unchanged with respect to wild type. Conversely, in *FC2*-overexpressing Arabidopsis plants, ALA synthesis rates decreased relative to wild type (Fig. 3d). As a result of their lower ALA synthesis rates, *p35SFC2* lines contained less Pchlide than the wild type, both in the light and the dark (Fig. 4b-c). In contrast to these consequences of deregulated FC2 levels, ALA synthesis rates were unchanged in leaves of both *fc1-1* mutants (Fan et al., 2019) and *FC1* overexpressor lines (Fig. S5).

These findings are consistent with the metabolic effects on TBS observed in *FC2-*deficient seedlings. Woodson et al. (2015) detected an accumulation of Proto in SD-grown Arabidopsis *fc2* knock-down seedlings at the beginning of the light phase. *FC2* antisense tobacco seedlings also contained higher Proto levels, which correspond with higher ALA synthesis rates relative to wild-type plants (Papenbrock et al., 2001). Moreover, it was reported that Arabidopsis *fc2* mutants grown under alternating light-dark conditions exhibit a *flu*-like phenotype owing to the accumulation of photoreactive Pchlide in the dark and photodynamic damage upon subsequent exposure to light (Scharfenberg et al., 2015). Based on our data, this Pchlide accumulation can be explained by the elevated rate of ALA synthesis in FC2-deficient seedlings, which also contain less POR.

We propose that in PORB- and FC2-deficient seedlings, POR is degraded, which in turn attenuates the efficacy of the FLU-dependent GluTR-inactivation complex. This metabolic perturbation leads to enhanced ALA synthesis (Fig. 3d) and thus to elevated Pchlide levels in dark-grown seedlings (Fig. 4b). Hence, the ALA synthesis-dependent increase in Pchlide levels is attributable to PORB deficiency and its need for adequate FLU-mediated inactivation of GluTR. Besides the elevated Pchlide levels, thanks to *FC2-*driven *FC1* expression in pFC2FC1(*fc2/fc2*), sufficient amounts of heme are available to prevent significant Proto accumulation, as shown in *FC2* knock-down mutants (Figs 2a and 4d; and Richter et al., 2019).

In contrast, *FC2* overexpression correlates with a slightly increased level of PORB, while the other enzymes of TBS analyzed, including GluTR, accumulate to wild-type levels (Fig. 3b). This, conversely, confirms that an FC2-mediated increase in POR stability potentiates GluTR inactivation and lowers ALA-synthesizing capacities (Fig 3d). It is worth mentioning here that PORB deficiency did not decrease FC2 levels (Figure 3b). These analyses emphasize the PORB-stabilizing role of FC2 in the feedback-regulated inactivation of ALA synthesis. This mechanism enables precise quantitative matching of TBS metabolites to the prevailing requirements, whether in the dark, under stress conditions or under fluctuating light intensities.

### Contributions of FC2 and POR to inactivation of ALA synthesis in concert with other regulatory mechanisms

Furthermore, the FC2-POR interaction points to a new function for FC2. Intriguingly, the POR-FC2 interaction involves enzymes that belong to different branches of TBS. We suggest that the interaction of these two proteins does not stimulate their activity, but rather results in their mutual inhibition. According to this regulatory concept, it seems sensible that synthesis of Mg-containing Chl and the FC2-catalyzed heme synthesis should be simultaneously feedback-controlled by the same mechanism of FLU-mediated inactivation of GluTR in darkness or adverse conditions. Since Chl is not synthesized in darkness, the demand for plastidic hemoproteins is also likely decreased.

It should be emphasized here that the involvement of FC2 in FLU-mediated suppression of ALA synthesis is another impact of plastidic heme synthesis on the post-translational feedback control of ALA synthesis, which originate from FC2-synthesized heme (Richter et al., 2019). The heme-dependent stimulation of GluTR degradation was discovered based on the promotion of GluTR degradation by binding of heme to GBP which attenuates the interaction of GBP with GluTR, and consequently exposes the latter to Clp-dependent proteolysis in dark-incubated Arabidopsis leaves.

With respect to the regulatory impact of the large set of factors on GluTR activity and stability (see Introduction), it is conceivable that the direct effect of de-regulated FC2 expression alters ALA synthesis only to some extent. The changes observed in ALA synthesis rate in the absence of FC2 and upon overproduction of the protein were about 60-70%. However, because of the crucial functions and the deleterious photodynamic actions of tetrapyrroles, plants have evolved multiple regulatory mechanisms and rely on their concerted action.

The model depicted in Fig. 8 illustrates the role of the two FC isoforms during day and night. Upon light exposure, chloroplasts activate extensive ALA synthesis which is initiated and controlled by GluTR. ALA synthesis is suggested to occur in the stroma, while a portion of GluTR is always found to be associated with the thylakoid membranes. It is assumed that GBP binds firmly to both stroma-localized and membrane-bound GluTR to prevent its degradation. While the major portion of TBS metabolites is directed into the Mg branch, both FC isoforms contribute to heme synthesis. Due to their distinct localization in plastids, it is proposed that FC1 delivers heme for all processes outside the plastids, and FC2 serves for plastid hemoproteins. In the dark, FC2 interacts with and stabilizes POR. Pchlide accumulation triggers the formation of the GluTR-inactivation complex. Membrane-bound FLU inhibits GluTR activity, leading to the suppression of ALA formation. Then, Chl and heme synthesis are simultaneously inactivated. It is assumed that a residual level of ALA-synthesizing activity in darkness channels metabolites to FC1 for the basal provision of heme mainly to proteins in the cytoplasm and mitochondria. It remains crucial to investigate the specific tasks of FC1 and the FC1-synthesized heme supply for the cellular metabolism, particular for the defense against stress conditions and retrograde signaling, independently of any possible potential contributions of FC2 in green leaves (Shimizu et al., 2019). In future, studies are needed to specify which characteristic structural traits of FC2 and FC1 are responsible for their specific tasks in heme synthesis and cellular regulation.

In conclusion, this report uncovers a novel contribution of FC2 to the post-translational control of TBS. Apart from its role in the dominant heme synthesis for plastid-localized proteins, FC2 participates in the control of dark-suppressed ALA synthesis. FC2 stabilizes POR and enhances the efficacy of the GluTR inactivation complex. Consequently, it is expected that FC2’s interaction with POR is likely to inactivate FC2-driven heme synthesis.

## ACKNOWLEDGEMENTS

This work was supported by the Chinese Scholarship Council to T.F. and the Deutsche Forschungsgemeinschaft to B.G (SFB TRR175, subproject C04). We thank Paul Hardy for critical reading of the manuscript.

## Supplemental Data

**Fig. S1**. Quantification of selected photosynthetic proteins in Col-0, pFC2FC1(fc2/fc2) and fc2-1 mutants.

**Fig. S2**. Biochemical analysis of the complemented fc2-2 lines (pFC2FC2(fc2/fc2)) and wild-type controls.

**Fig. S3**. Interaction analyses of FC and CHL27 and the two POR isoforms PORA and PORC by BiFCs.

**Fig. S4**. Interaction of FC 2 and PPO 1 by BiFC.

**Fig. S5**. ALA synthesis rates in FC1 overexpression lines. Table S1 List of primers used in this study

