## Supplemental information for "FC2 stabilizes POR and suppresses ALA formation in the tetrapyrrole biosynthesis pathway"

**Supplemental Figures**

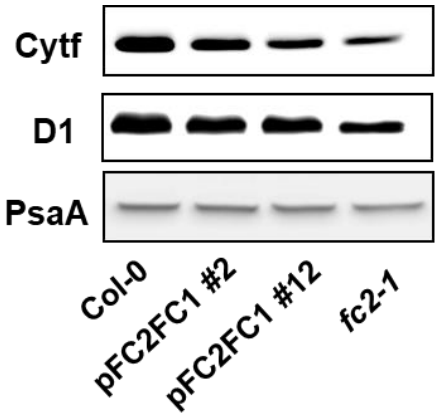


**Fig. S1 Quantification of selected photosynthetic proteins in Col-0, pFC2FC1(*fc2/fc2*) and *fc2-1* mutants**.

Total protein extracts of leaves from 5-week-old SD seedlings of the indicated lines were fractionated by SDS-PAGE. After blotting, selected proteins were analysed with specific antibodies: Cytf, cytochrome f, a heme-binding protein of the cytochrome b6f complex, D1: component of PSII core complex; PsaA; a main subunit of the PSI core complex.

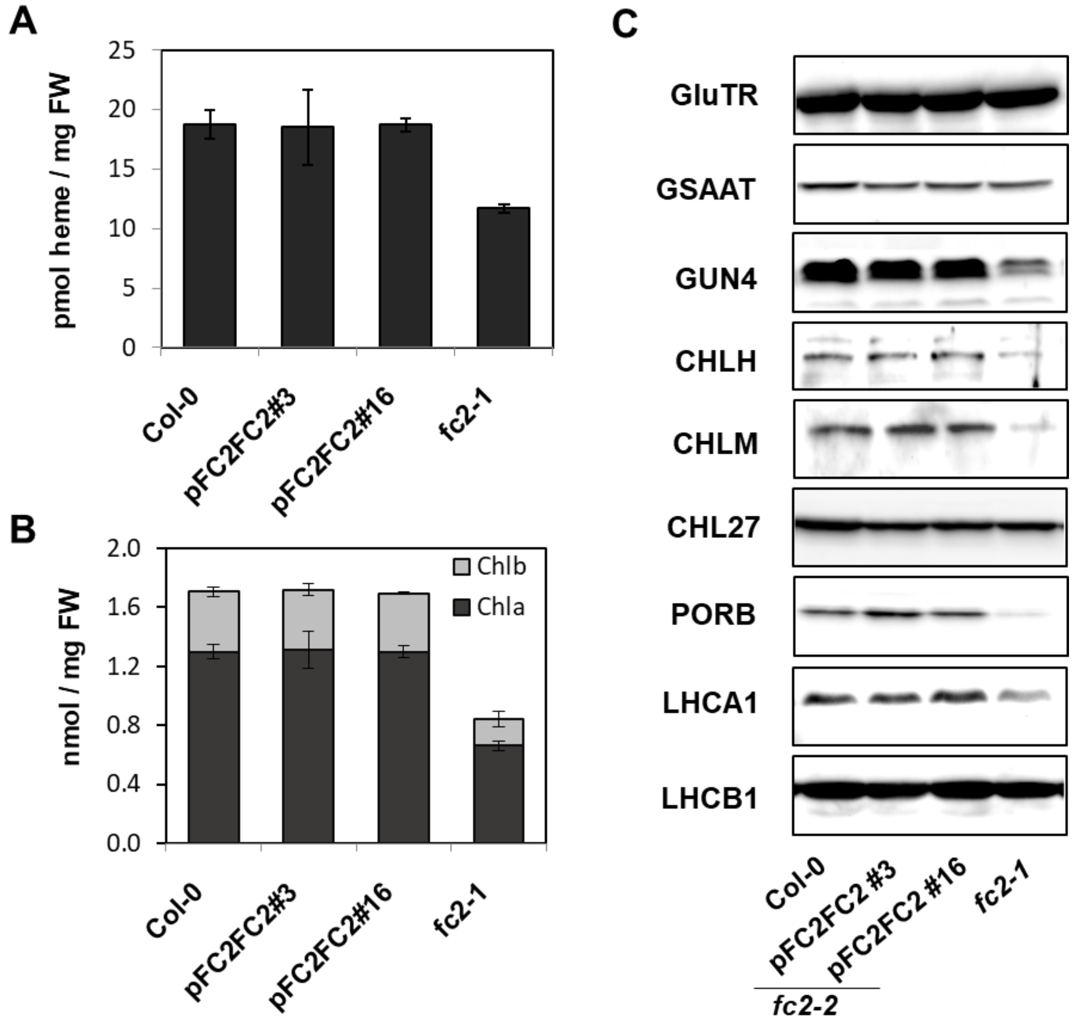


**Fig. S2 Biochemical analysis of complemented *fc2-2 lines* (pFC2FC2(*fc2/fc2*)) and wild-type controls.**

(**a**,**b**) Quantification of end products of the TBS pathway. Heme (**A**) and chlorophyll (**B**) were extracted from leaf samples of 3-week-old, SD-grown seedlings of pFC2FC2(*fc2/fc2*), *fc2-1* and wild-type (Col-0) plants. (**c**) Analyses of selected proteins from SD-grown plants. GluTR: glutamyl-tRNA reductase; GSAAT*:* glutamate-1-semialdehyde aminotransferase; GUN4: GENOMES UNCOUPLED 4; CHLH: Mg chelatase subunit H; CHLM: Mg-protoporphyrin IX methyltransferase; PORB: protochlorophyllide oxidoreductases B; LHCA1/LHCB1: light-harvesting chlorophyll-binding proteins A1 and B1.

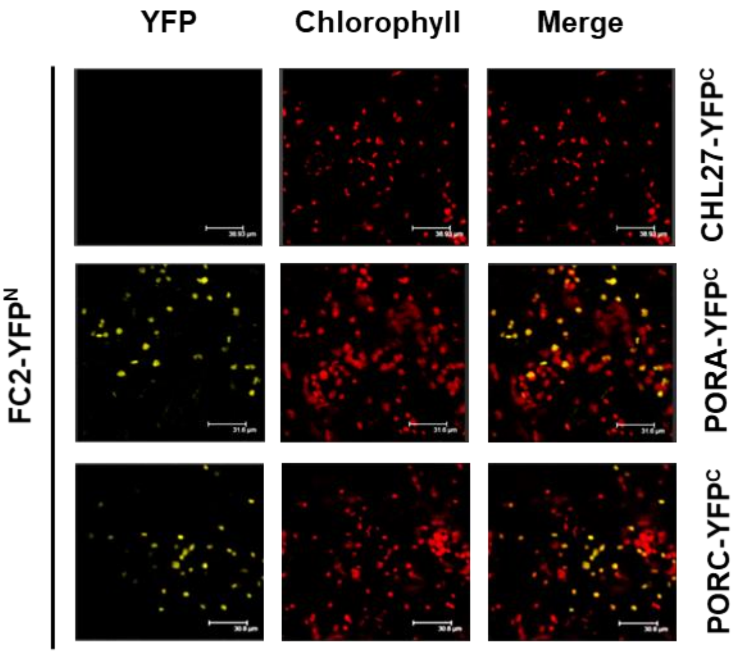


**Fig. S3 Interaction analyses of FC2 with CHL27, PORA and PORC by BiFC.**

Protein-protein interaction in chloroplasts was visualized by BiFC assays. YFP images show tobacco (*Nicotiana benthamiana*) cells transiently expressing gene constructs encoding the fusion proteins indicated. The red fluorescence (Chlorophyll) represents chloroplast autofluorescence, and overlay images (Merge) show merged signals. The fluorescence patterns shown are typical of those seen in more than three independent replicates. Scale bars were indicated.

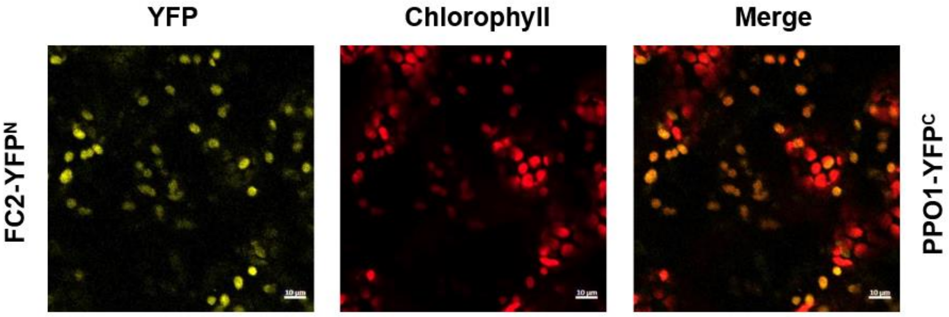


**Fig. S4 Interaction analyis of FC2 and PPO1 using BiFC.**

Protein-protein interaction in chloroplasts was visualized by BiFC assay. YFP images show tobacco (*Nicotiana benthamiana*) cells transiently expressing gene constructs encoding the fusion proteins indicated. The red fluorescence represents chloroplast autofluorescence, and overlay images show merged signals from both types of fluorescence. The images are typical of the data obtained in more than three replicates. The cYFP-tagged PPOX1 was co-expressed with nYFP-tagged FC2 in tobacco leaf cells. Scale bars are indicated.

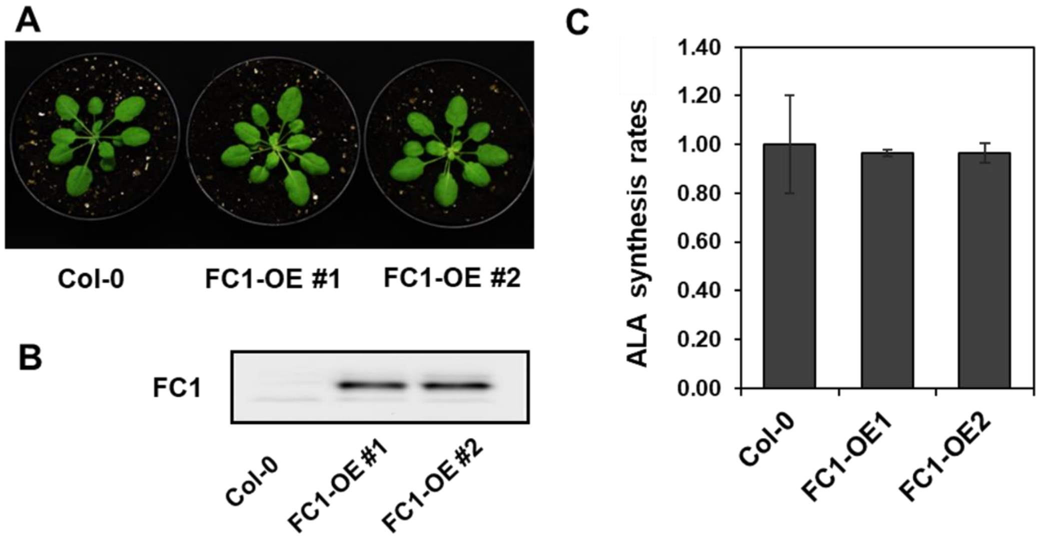


**Fig. S5 ALA synthesis rates in FC1 overexpression lines.**

(**A**) Growth of FC1 overexpression (FC1-OE) lines and wild-type seedlings under SD conditions.

(**B**) FC1 protein levels in leaf extracts obtained from leaves of the seedlings shown in panel A.

(**C**) Relative ALA synthesis rates of 3-week-old FC1-OE mutants and wild-type seedlings. Error bars indicate standard deviations (n≥3).

**Table S1 List of primers used in this study**

| | Forward (Fw, 5'-3') | Reverse (Rev, 5'-3') |
|---|---|---|
| **qRT-PCR** | | |
| HEMA1 | TTGCTGCCAACAAAGAAGAC | CCGTCTCCAATGAATCCCTC |
| FLU | AAGCCATACAGTATCACTCCA | TCCAGAATCTTCACTTTCCCT |
| GBP | CAGTTGACCGTGTTCTCC | AATCCAAGCCTATCCATC |
| CHLD | GGGAAGATATACAAGGCAGGG | GAGATTACAGCATCCGAAGCA |
| GUN4 | TTAGGACACTTACCGCTCAC | GCTCGTCTTCTGTTTCTCCA |
| CHLM | TTGCTGAAGCTGAGATGAAGGCAAAG | CAACGGTATCATACTTCCCAGTTAGG |
| PORA | TACCCTCTTCCCTCCTTTCC | GTTCCAGCTCCAATACACTCC |
| PORB | TGATTACCCTTCAAAGCGTCTCA | CAATGTATTCGTGTTCCCGGT |
| PORC | CTGGCAAAAGACTAGCACAGGTT | CAATACACTCCTGACTTCCCAAGAC |
| HO1 | GCGGATACTCGATAATAAAGAACTC | TCTTCTCTAGTCCACTCCTCTG |
| HO2 | GCTGGAGTTCAATAGGTGGG | TTAAATGCCTTTGCCGTTTCC |
| ACTIN | CTTCCCTCAGCACATTCCAG | GACCTGCCTCATCATACTCG |
| **BiFC constructs** | | |
| FC1 | CAAAAAAGCAGGCTGAGCTATGCAGGCAACGG | CAAGAAAGCTGGGTGTAGGTTCCGGAACGCATG |
| FC2 | CAAAAAAGCAGGCTGAGCAATGAATTGCCCAGC | CAAGAAAGCTGGGTGTAATGAAGGCAAGATGCC |
| **Yeast constructs** | | |
| pNub-FC1 | CAAAAAAGCAGGCTTATGCGATATAAAAGAGAGATCTTTCG | CAAGAAAGCTGGGTGTCATAGGTTCCGGAACGCATGGAACATC |
| pDHB-FC1 | CACCTAGGTGCGATATAAAAGAGAGATCTTTCG | CACTGCAGTAGGTTCCGGAACGCATGGAACATC |
| pNub-FC2 | CAAAAAAGCAGGCTTAGCATTTGCTGCTACTTCATCAAAC | CAAGAAAGCTGGGTGTCATAATGAAGGCAAGATGCCCCACTG |
| pDHB-FC2 | CACCTAGGGCATTTGCTGCTACTTCATCAAAC | CACTGCAGTAATGAAGGCAAGATGCCCCACTG |
| **Generation of transgenic plants** | | |
| pFC2FC1 (fc2/fc2) | pFC2_fw ATGATATCGATACATTTGAAGACAATGC | pFC2_rev ATGATATCGTTTAAACGAAACTTGAAGAAACCTTAAC |
| | FC1_TPfw ACACGTGCATGGTGTGGTCCCCTTAAT | FC1_stop ACACGTGCTATAGGTTCCGGAACGC |
| pFC2FC2 (fc2/fc2) | pFC2_fw ATGATATCGATACATTTGAAGACAATGC | FC2_stop ATCACGTGATTATAATGAAGGCAAGATGCC |
| 35SFC2 | FC2_TPfw ACACGTGAAAATGAATTGCCCAGCCATG | FC2_stop ATCACGTGATTATAATGAAGGCAAGATGCC |
